# Calcineurin inhibition deactivates pyruvate dehydrogenase and induces proximal tubule cell metabolic dysfunction, causing profibrotic phenotype

**DOI:** 10.1101/2024.11.20.624584

**Authors:** Yasuhiro Oda, Hiroshi Nishi, Fumie Hamano, Teruhiko Yoshida, Yoshihiro Kita, Jeffrey B. Kopp, Masaomi Nangaku

**Author notes:** **Address for correspondence:** Hiroshi Nishi, M.D., Ph.D., Division of Nephrology and Endocrinology, The University of Tokyo Graduate School of Medicine 7-3-1 Hongo, Bunkyo-ku, Tokyo 113-8655 Japan.

## Abstract

Calcineurin inhibitors (CNIs) are indispensable immunosuppressants for transplant recipients and patients with autoimmune diseases, but chronic use causes nephrotoxicity, including kidney fibrosis. Why inhibiting calcineurin, a serine/threonine phosphatase, causes kidney fibrosis remains unknown. We performed single-nucleus RNA sequencing of the kidney from a chronic CNI nephrotoxicity mouse model and found an increased proportion of injured proximal tubule cells, which exhibited altered expression of genes associated with oxidative phosphorylation, cellular senescence and fibrosis. In cultured primary human renal proximal tubule epithelial cells, CNIs caused phosphorylation (deactivation) of pyruvate dehydrogenase, impaired mitochondrial metabolism and senescence-associated phenotypes, all of which were ameliorated by pyruvate dehydrogenase activation. Finally, administration of dichloroacetic acid, a known activator of pyruvate dehydrogenase, in the chronic CNI nephrotoxicity mouse model mitigated kidney fibrosis and the associated transcriptional changes. Collectively, calcineurin inhibition deactivates pyruvate dehydrogenase and induces proximal tubule cell metabolic dysfunction, causing profibrotic phenotype.

## INTRODUCTION

Calcineurin inhibitors (CNIs) are immunosuppressive drugs that are indispensable for patients after organ or hematopoietic transplantation and for patients with autoimmune disorders. Cyclosporin A (CsA) and tacrolimus, also known as FK506, are the two CNIs that have been used for these purposes for decades. Despite their substantial efficacy in immunosuppression, chronic nephrotoxicity has been a major adverse effect associated with the use of CNI since early clinical studies in the 1980s^1,2^ and remains a serious issue.^3,4^ There is no clinically available preventive strategy for CNI nephrotoxicity, leaving clinicians with the need to balance the benefit of immunosuppression against the risk of chronic nephrotoxicity, by adjusting the dosage of CNI. Insufficient immunosuppression causes allograft rejection, whereas higher doses of CNI increase the risk of advanced kidney disease requiring lifelong kidney replacement therapy. Both outcomes have deleterious impacts on the lives of patients.

While reversible, acute CNI nephrotoxicity is caused by vasoconstriction of glomerular afferent arterioles and subsequent reduction in renal blood flow,^5,6^ the mechanism underlying irreversible, chronic CNI nephrotoxicity has not been fully elucidated.^3,4^ The hallmarks of chronic CNI nephrotoxicity include kidney fibrosis, which is an irreversible chronic lesion and the central cause of kidney dysfunction.^3,4^ The current concept of the etiology of chronic CNI nephrotoxicity is that vascular injury followed by arterial hyalinization and subsequent glomerular hypoperfusion contributes to the development of kidney fibrosis. However, this pathologic pathway has not been shown directly^4^ and remains a theory based on morphological findings.

Meanwhile, limited attention has been directed to the physiological role of calcineurin in normal kidneys and on whether its inhibition can cause kidney fibrosis. Calcineurin is a Ca^2+^-calmodulin-dependent serine/threonine phosphatase conserved from fungi to humans.^7,8^ Previous studies have shown that regulation of cellular metabolism is one of the many functions of calcineurin.^9–11^ A recent study demonstrated that calcineurin dephosphorylates and thereby activates pyruvate dehydrogenase, which is a gatekeeper that fuels the tricarboxylic acid (TCA) cycle and controls subsequent oxidative phosphorylation in mitochondria, as shown in an immortalized human cell line.^11^

Here, we show that 1) calcineurin inhibition deactivates pyruvate dehydrogenase and impairs mitochondrial energy metabolism in energy-demanding proximal tubule cells in the kidneys and that 2) the impaired mitochondrial metabolism causes a proinflammatory, profibrotic phenotype associated with cellular senescence, thereby contributing to kidney fibrosis. We performed a comprehensive analysis of transcriptomic changes in the early stage of chronic CNI nephrotoxicity mouse model at single-cell resolution and found an increased proportion of injured proximal tubule cells. These cells exhibited altered expression of genes associated with oxidative phosphorylation, cellular senescence and fibrosis. Analyses of publicly available transcriptomes of human kidneys confirmed that some of these manifestations are observed in human allograft kidneys under certain conditions. *In vitro* experiments with primary human renal proximal tubule epithelial cells (RPTECs) revealed that CNIs cause phosphorylation (deactivation) of pyruvate dehydrogenase, impaired mitochondrial metabolism and senescence-associated phenotypes, all of which were ameliorated by pyruvate dehydrogenase activation. Finally, administration of dichloroacetic acid (DCA), an activator of pyruvate dehydrogenase, in the chronic CNI nephrotoxicity mouse model mitigated kidney fibrosis and decreased the expression of genes associated with proximal tubule injury, cellular senescence and fibrosis. These results provide a novel perspective in understanding chronic CNI nephrotoxicity and suggest a therapeutic approach.

## RESULTS

### KIM-1–positive injured proximal tubule cell proportion is increased in chronic CNI nephrotoxicity mouse model from an early stage

A chronic CNI nephrotoxicity mouse model was created by administering CsA as previously described^12^ with some modifications. After 4 weeks of administering CsA, mice developed kidney fibrosis (Figure 1A) and mild kidney dysfunction (Figure 1B).

**Figure 1.**
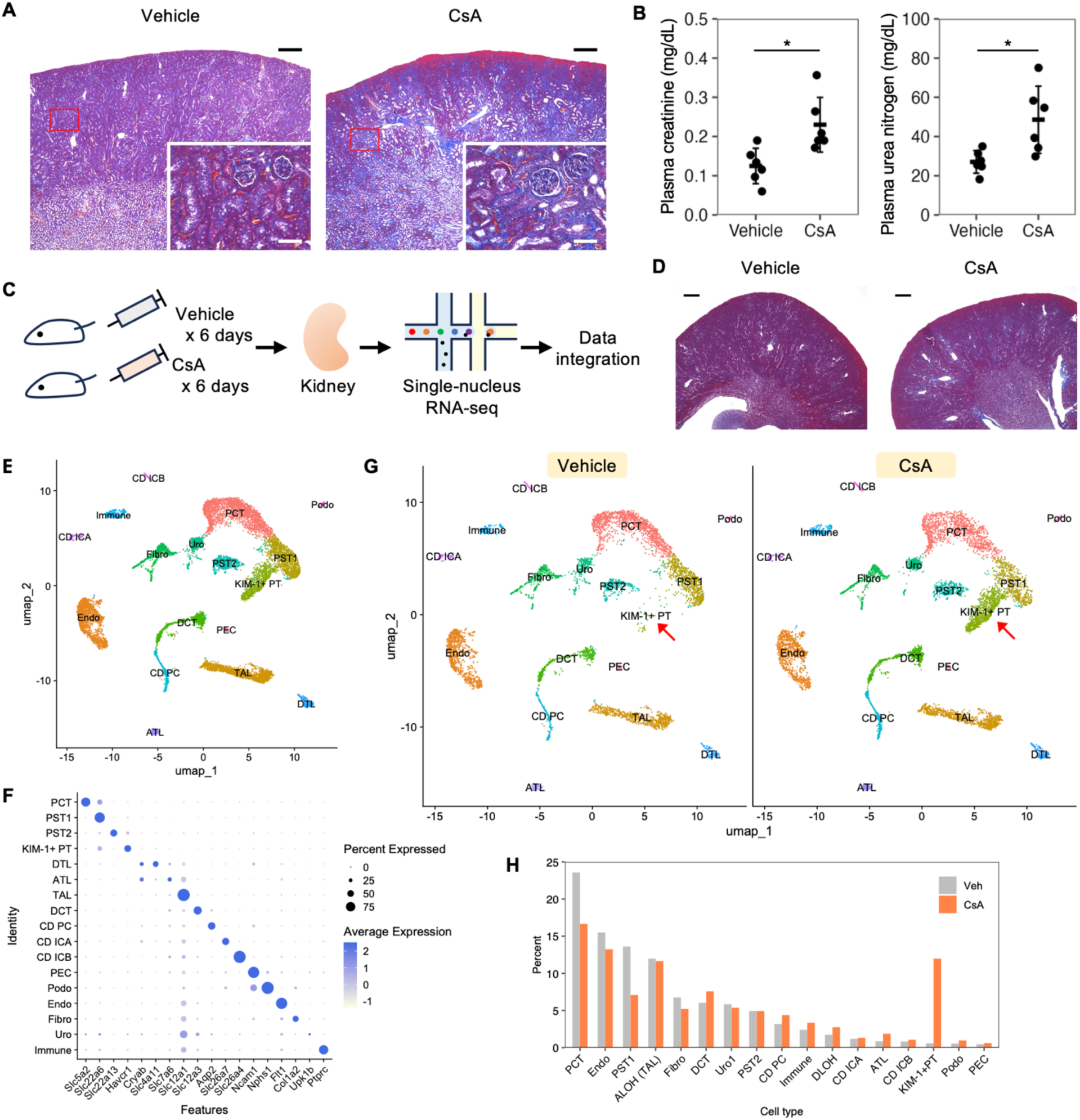
KIM-1–positive injured proximal tubule cell proportion is increased in chronic CNI nephrotoxicity mouse model from an early stage. (A) Masson’s trichrome staining on kidney from ICR mice receiving vehicle or cyclosporin A (CsA) for 4 weeks. High-power field images show the area in red rectangles. Scale bar, 300 μm for low-power field and 30 μm for high-power field images. (B) Plasma creatinine and urea nitrogen levels in ICR mice receiving vehicle or CsA for 4 weeks (n = 6 per group). (C) Experimental design of single-nucleus RNA sequencing of the kidneys from early-stage chronic calcineurin inhibitor nephrotoxicity mouse model and its control. (D) Masson’s trichrome histology staining on kidney from ICR mice receiving vehicle or CsA for 6 days. Scale bar, 300 μm. (E) UMAP presentation of 12,695 cells profiled from the kidney of early-stage chronic calcineurin inhibitor nephrotoxicity mouse model and its control. Doublets are removed. PCT, proximal convoluted tubule; PST, proximal straight tubule; KIM-1+ PT, KIM-1–positive proximal tubule; DTL, descending thin limb of Henle’s loop; ATL, ascending thin limb of Henle’s loop; TAL, thick ascending limb of Henle’s loop; DCT, distal convoluted tubule; CD PC, collecting duct – principal cell; CD ICA, collecting duct – type A intercalated cell; CD ICB, collecting duct – type B intercalated cell; PEC, parietal epithelial cell; Podo, podocyte; Endo, endothelial cell; Fibro, fibroblast; Uro, uroepithelial cell; Immune, immune cell. (F) Dot plot showing the expression pattern of cluster-specific marker genes. Doublets are removed. (G) UMAP presentation of cells profiled from the kidney of early-stage chronic calcineurin inhibitor nephrotoxicity mouse model (7,188 cells) or its control (5,507 cells). Red arrow designates KIM-1– positive proximal tubule cells. (H) Bar chart showing the proportion of each cell type in each sample. Data are presented as mean ± SD. Unpaired two-tailed Welch’s t test was used. n represents the number of biological replicates. * *p* < 0.05. See also Figure S1–S2.

To characterize the transcriptional changes in kidney cells in an early-stage chronic CNI nephrotoxicity mouse model at single-cell resolution, kidneys from mice that received either CsA or its vehicle for 6 days were subjected to single-nucleus RNA sequencing (snRNA-seq; Figure 1C). Kidney fibrosis was negligible at this early stage (Figure 1D). Unsupervised clustering of snRNA-seq data (Figure S1; See also Methods) and removal of doublets identified 17 clusters of cells encompassing all major cell types in the kidneys (Figure 1E). Cell types were annotated using lineage markers, as shown in Figure 1F. The proportion of injured proximal tubule cells, characterized by the upregulation of the *Havcr1* gene (coding for kidney injury molecule-1 [KIM-1]), was substantially higher in the early-stage chronic CNI nephrotoxicity mouse model than in its control (Figures 1G, 1H and S2A). A portion of these KIM-1–positive injured proximal tubule cells also expressed the *Vcam1* gene (coding for vascular cell adhesion molecule 1 [VCAM-1]),^13,14^ which designates failed-repair proximal tubule cells (Figure S2B).^14^ These results show that CsA administration increased the proportion of KIM-1–positive injured proximal tubule cells, including failed-repair proximal tubule cells, in the chronic CNI nephrotoxicity mouse model at an early stage before the development of kidney fibrosis.

### KIM-1–positive injured proximal tubule cells show transcriptional changes in genes associated with oxidative phosphorylation, cellular senescence and fibrosis

KIM-1–positive injured proximal tubule cells and VCAM-1–positive failed-repair proximal tubule cells are known to exhibit a proinflammatory and profibrotic phenotype, as well as the senescence-associated secretory phenotype (SASP).^14–18^ Increasing evidence has demonstrated that injured proximal tubule cells contribute to the progression of chronic kidney disease (CKD) of various etiologies.^14–18^ Mitochondrial dysfunction is a common cause of proximal tubule cell injury and a major cause of cellular senescence accompanied by secretion of proinflammatory, profibrotic SASP factors.^19–21^

To characterize the transcriptional changes in KIM-1–positive injured proximal tubule cells in the chronic CNI nephrotoxicity mouse model, we next performed gene set enrichment analysis (GSEA) to compare differentially expressed genes (DEG) among cell clusters. A combined cluster of proximal tubule cells—namely, proximal convoluted tubule cells, proximal straight tubule cells, and KIM-1–positive injured proximal tubule cells—in early-stage chronic CNI nephrotoxicity mouse model showed down-regulated expression of genes associated with oxidative phosphorylation (false discovery rate [FDR] < 1×10^−15^) and up-regulated expression of genes associated with nuclear factor kappa B (NF-κB) signaling pathway (FDR < 1×10^−15^) compared to proximal tubule cells in the control (Figure 2A). We further compared KIM-1– positive injured proximal tubule cells to other proximal tubule cells in the early-stage chronic CNI nephrotoxicity mouse model. KIM-1–positive injured proximal tubule cells showed a lower expression of genes associated with oxidative phosphorylation (FDR < 1×10^−15^) and a higher expression of genes associated with NF-κB signaling pathway (FDR < 1×10^−15^) compared to other proximal tubule cells in the early-stage chronic CNI nephrotoxicity mouse model (Figure 2B). NF-κB signaling is one of the proinflammatory signaling pathways that regulate the expression of SASP factors.^22^

**Figure 2.**
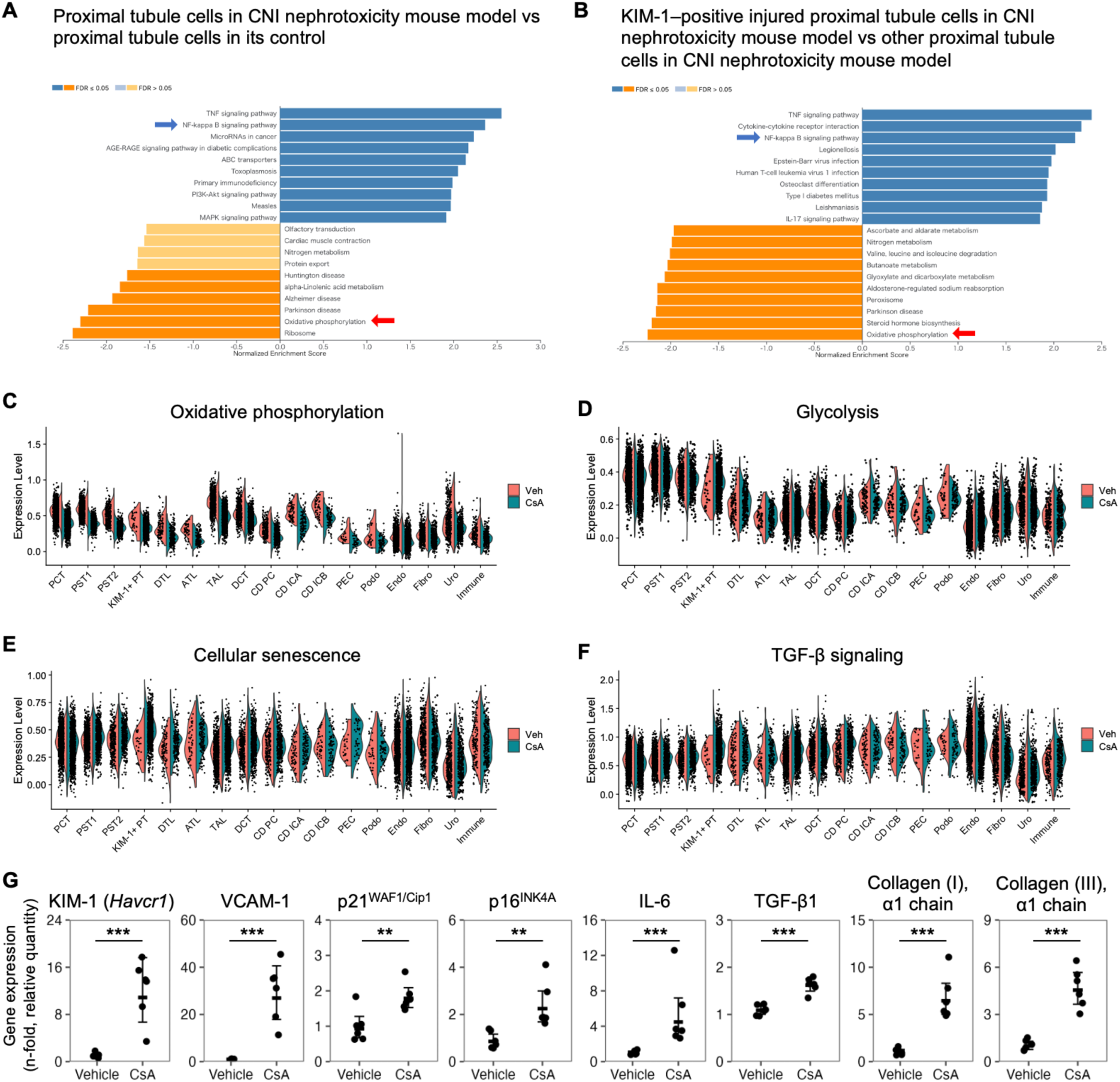
KIM-1–positive injured proximal tubule cells show transcriptional changes in genes associated with oxidative phosphorylation, cellular senescence and fibrosis. (A and B) Bar chart showing top KEGG pathways enriched in proximal tubule cells in calcineurin inhibitor (CNI) nephrotoxicity mouse model vs proximal tubule cells in its control (A) and in KIM-1– positive injured proximal tubule cells in CNI nephrotoxicity mouse model vs other proximal tubule cells in the CNI nephrotoxicity mouse model (B). Blue bars show enrichment in the experimental group, whereas orange bars show enrichment in the control group. Dark color designates FDR ≤ 0.05, and light color designates FDR > 0.05. Blue arrow points to NF-κB signaling pathway, and red arrow points to oxidative phosphorylation. FDR, false discovery rate. (C–F) Violin plot showing the enrichment of genes associated with oxidative phosphorylation (C), glycolysis (D), cellular senescence (E) and TGF-β signaling (F). (G) mRNA expression of genes associated with proximal tubule injury, cellular senescence and fibrosis measured by quantitative PCR in the kidneys of ICR mice receiving vehicle or cyclosporin A (CsA) for 2 weeks (n = 6 per group). Data are presented as mean ± 1.96 SE. Unpaired two-tailed Welch’s t test was used. n represents the number of biological replicates. ** *p* < 0.01, *** *p* < 0.001. See also Figure S3.

We next compared the expression levels of genes associated with oxidative phosphorylation, cellular senescence and transforming growth factor beta (TGF-β) signaling in different cell types. The expression of genes associated with oxidative phosphorylation was markedly down-regulated in proximal tubule cells as well as in other tubule cells, and this downregulation was more prominent than that in endothelial cells (Figure 2C). Downregulation of expression was not common to all metabolic pathways. For example, the expression of genes associated with glycolysis did not show a remarkable change between the early-stage chronic CNI nephrotoxicity mouse model and its control in all cell types (Figure 2D). Moreover, expression of genes associated with cellular senescence and TGF-β signaling was up-regulated in KIM-1–positive injured proximal tubule cells in the early-stage chronic CNI nephrotoxicity mouse model compared to other tubular cells (Figures 2E and 2F). Uniform manifold approximation and projection (UMAP) plots of the expression levels of these gene sets are illustrated in Figure S3.

To confirm the transcriptional changes induced by administering CsA in a larger sample size, kidneys from mice after administration of vehicle or CsA for 14 days were subjected to RNA extraction and quantitative PCR. Marker genes for identifying injured proximal tubules (KIM-1 and VCAM-1), cellular senescence (p21^WAF1/Cip1^, p16^INK4A^, interleukin [IL]-6 and TGF-β1) and fibrosis (TGF-β1, α1 chain of type I collagen and α1 chain of type III collagen) were all up-regulated in the kidneys of early-stage chronic CNI nephrotoxicity mouse model (Figure 2G).

Taken together, these observations indicate that CsA increases the number of KIM-1– positive injured proximal tubule cells with down-regulated expression of genes associated with oxidative phosphorylation and up-regulated expression of genes related to cellular senescence and fibrosis.

### Transcriptome of human allograft kidneys under certain conditions shows down-regulated expression of genes associated with oxidative phosphorylation

To investigate whether these transcriptional changes are observed in human kidneys after CNI exposure, we analyzed publicly available transcriptome datasets. While we found no available dataset that precisely compared the transcriptomes of normal human kidneys with those after CNI exposure, we did identify two datasets that were suitable to answer the question.

The first resource (Gene Expression Omnibus [GEO] accession number GSE53605) included microarray datasets of normal kidney allografts (n = 18) and kidney allografts with CNI nephrotoxicity diagnosed by histological evidence (n = 14).^23^ GSEA of the DEG between the two groups revealed that genes associated with oxidative phosphorylation (FDR < 1×10^−15^) and TCA cycle (FDR < 1×10^−15^) were among the most down-regulated gene sets in kidney allografts with CNI nephrotoxicity (Figure 3A). In contrast, genes associated with the NF-κB signaling pathway was up-regulated in these kidney allografts (FDR < 1×10^−15^, Figure 3A). These findings are consistent with our observations in the experimental mouse model shown in Figure 2. Nevertheless, this analysis of human kidney transcriptomes might be merely observing a transcriptional difference between normal kidneys and the state of CKD, because the estimated glomerular filtration rate of the CNI group (35.3 ± 7.9 mL/min/1.73 m^2^) was remarkably lower than that of the normal kidney group (71 ± 11.3 mL/min/1.73 m^2^).^23^

**Figure 3.**
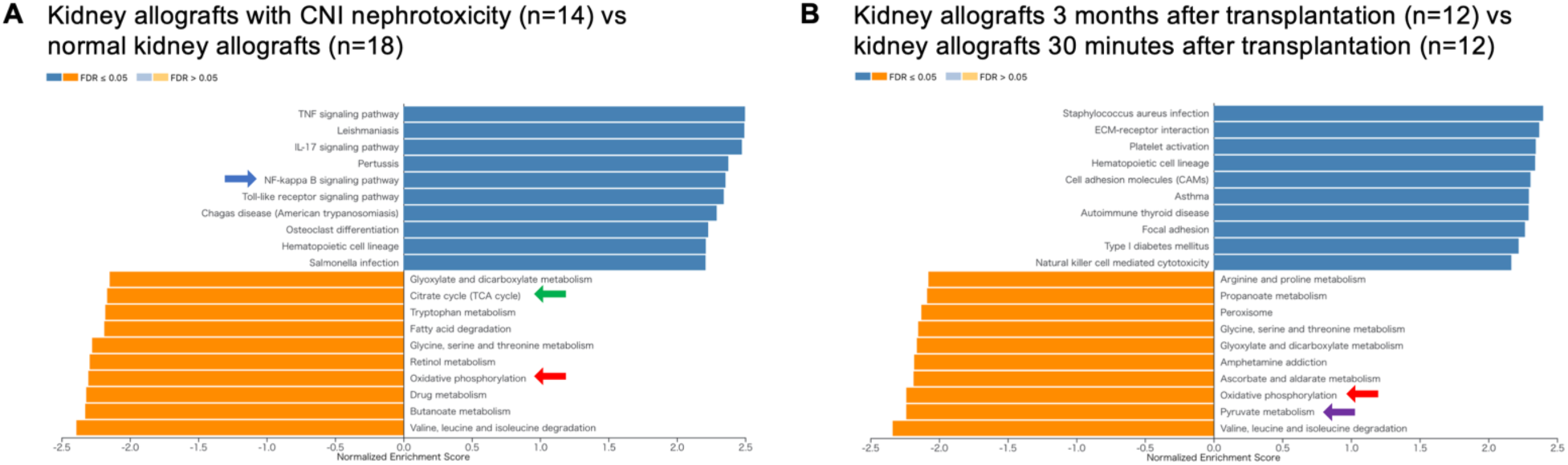
Transcriptome of human allograft kidneys under certain conditions shows down-regulated expression of genes associated with oxidative phosphorylation. (A and B) Bar chart showing top KEGG pathways enriched in kidney allografts with calcineurin inhibitor (CNI) nephrotoxicity vs normal kidney allografts (A) and in kidney allografts 3 months after transplantation vs kidney allografts 30 minutes after transplantation (B). Blue bars show enrichment in the sample group, whereas orange bars show enrichment in the control group. Dark color designates FDR ≤ 0.05, and light color designates FDR > 0.05. Blue arrow points to NF-κB signaling pathway, red arrow points to oxidative phosphorylation, green arrow points to citrate cycle (TCA cycle), and purple arrow points to pyruvate metabolism.

Therefore, we next analyzed the second resource (GEO accession number GSE178689), which comprises microarray datasets of kidney allograft biopsy specimens obtained 30 minutes (n = 12) and 3 months (n = 12) post-transplantation from the same patients.^24^ As the authors discuss, the latter samples are considered to represent changes in the kidneys with CNI exposure, adaptive immunity, and repair from perioperative ischemic injury.^24^ Interestingly, GSEA showed that genes associated with oxidative phosphorylation (FDR < 1×10^−15^) and pyruvate metabolism (FDR < 1×10^−15^) were among the most down-regulated gene sets in kidney allografts 3 months post-transplantation when compared to the same allografts 30 minutes post-transplantation (Figure 3B).

Although these analyses are not a direct comparison between the presence and absence of CNI exposure, these real-world data illustrate that kidney allografts after the perioperative period of transplantation and kidney allografts with CNI nephrotoxicity have lower expression of genes associated with oxidative phosphorylation compared to their counterparts.

### CNIs cause pyruvate dehydrogenase deactivation and metabolic dysfunction in primary human RPTECs

To characterize phenotypic changes that CNIs cause in proximal tubule cells, primary human RPTECs were cultured with 10 μM CsA, 10 μM tacrolimus or control (0.1% v/v dimethyl sulfoxide) for 24 h. In primary human RPTECs exposed to CNIs, the amount of phosphorylated, inactivated pyruvate dehydrogenase was increased (Figures 4A and 4B), and pyruvate dehydrogenase activity was decreased (Figure 4C). These results are consistent with a previous report that showed the direct activation of pyruvate dehydrogenase by calcineurin.^11^

**Figure 4.**
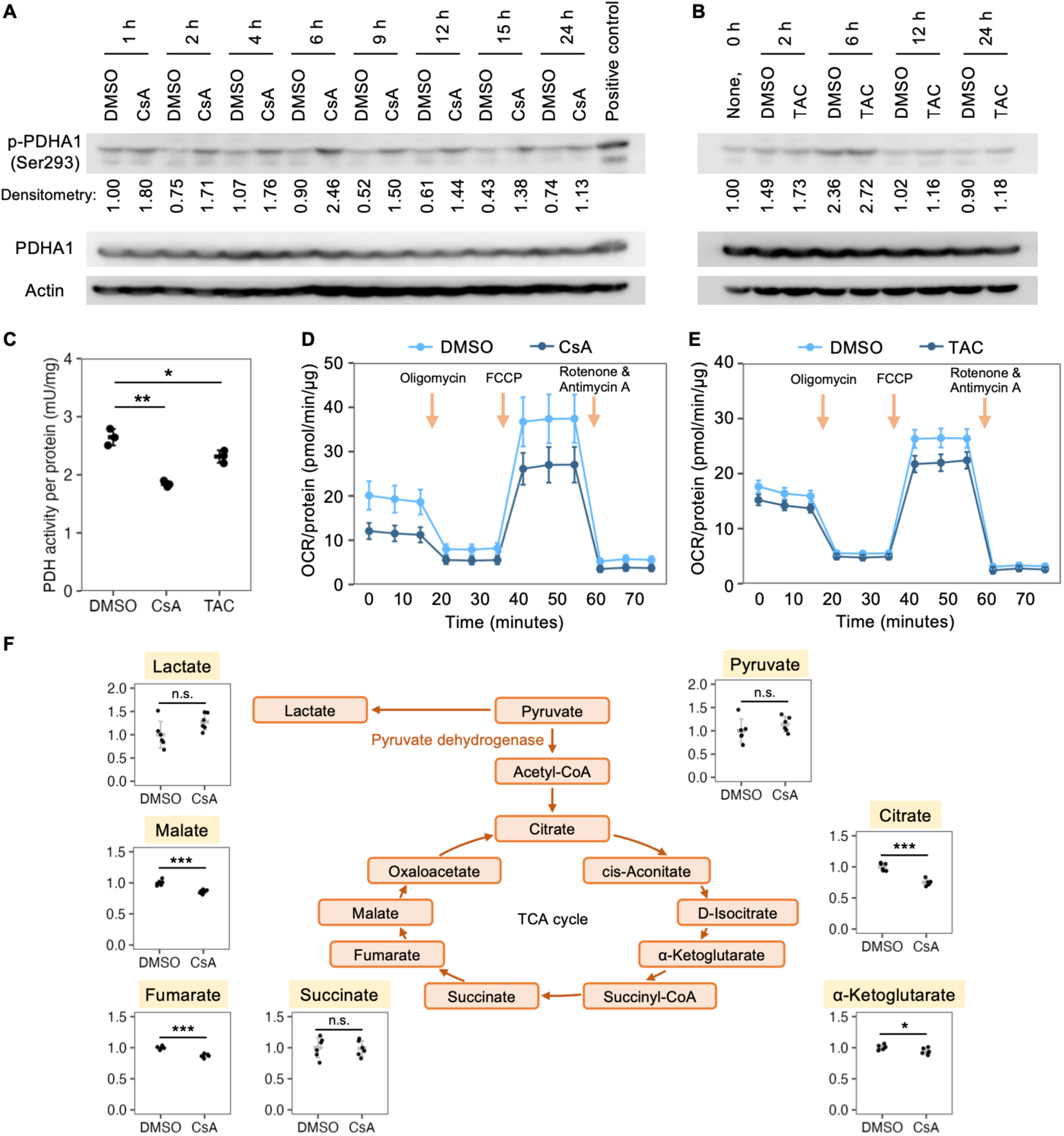
Calcineurin inhibitors cause pyruvate dehydrogenase deactivation and metabolic dysfunction in primary human renal proximal tubule epithelial cells. (A and B) Western blot of PDHA1 phosphorylation at Ser293, total PDHA1 and actin in primary human RPTECs cultured with DMSO or CsA (A) or with DMSO or TAC (B) for different length of time. Positive control is a sample known for containing phosphorylated PDHA1. CsA, cyclosporin A; DMSO, dimethyl sulfoxide; PDHA1, pyruvate dehydrogenase E1 subunit alpha 1; RPTEC, renal proximal tubule epithelial cell; TAC, tacrolimus. (C) Pyruvate dehydrogenase activity normalized to total protein levels in primary human RPTECs cultured with DMSO, CsA or TAC for 24 h (n = 3 per group). (D and E) Oxygen consumption rate (OCR) normalized to total protein levels in primary human RPTECs cultured with DMSO or CsA (D) or with DMSO or TAC (E) for 24 h (n = 12–15 per group). FCCP, carbonyl cyanide 4-(trifluoromethoxy)phenylhydrazone. (F) Mass spectrometric quantification (relative amount) of intracellular metabolites related to TCA cycle in primary human RPTECs cultured with DMSO or CsA for 24 h (n = 6). Data are presented as mean ± SD (C and F) or mean ± 1.96 SE (D and E). Unpaired two-tailed Welch’s t test was used. Holm’s correction for multiple comparisons was used in (C). n represents the number of biological replicates. * *p* < 0.05, ** *p* < 0.01, *** *p* < 0.001. n.s., not significant.

To assess whether cellular metabolism was altered by adding CNIs to the culture medium, we examined the level of mitochondrial respiration and the amount of intracellular TCA cycle metabolites. Extracellular flux analysis showed that the mitochondrial oxygen consumption rate was decreased in primary human RPTECs cultured with CNIs (Figures 4D and 4E) as previously reported.^25^ Gas chromatography–mass spectrometry revealed that primary human RPTECs cultured with CsA for 24 h contained lower amount of citrate, α-ketoglutarate, fumarate and malate, all of which are TCA cycle metabolites downstream of pyruvate dehydrogenase (Figure 4F). In contrast, the amounts of lactate and pyruvate, which are upstream of pyruvate dehydrogenase, were not decreased (Figure 4F). The average amounts of lactate and pyruvate in the samples were higher in the cells cultured with CsA; however, the difference was not statistically significant (lactate, *p* = 0.08; pyruvate, *p* = 0.28).

These observations demonstrate that a CNI causes deactivation of pyruvate dehydrogenase and metabolic dysfunction in primary human RPTECs.

### CNIs cause proinflammatory and profibrotic phenotypes associated with cellular senescence in primary human RPTECs

Metabolic dysfunction is a major trigger of cellular senescence, where SASP may cause proinflammatory and profibrotic responses and promote organ fibrosis.^19–21^ Given that CNIs impair cellular metabolism in primary human RPTECs, we next examined whether CNIs cause proinflammatory and profibrotic phenotypes associated with cellular senescence. Primary human RPTECs cultured with CNIs showed up-regulated expression of genes associated with cellular senescence (Figures 5A and S4), including *CDKN1A* (coding for p21^WAF1/Cip1^), *CDKN2A* (tested for mRNA variants coding for p16^INK4A^), *IL1B* (coding for IL-1β), and *TGFB1* (coding for TGF-β1). An increase in the protein expression of p21^WAF1/Cip1^ was confirmed after 5 days of incubation (Figure 5B). Furthermore, elevated activity of senescence-associated β-galactosidase was detected (Figures 5C and 5D). Thus, primary human RPTECs exhibit proinflammatory and profibrotic phenotypes associated with cellular senescence after incubation with CNI.

**Figure 5.**
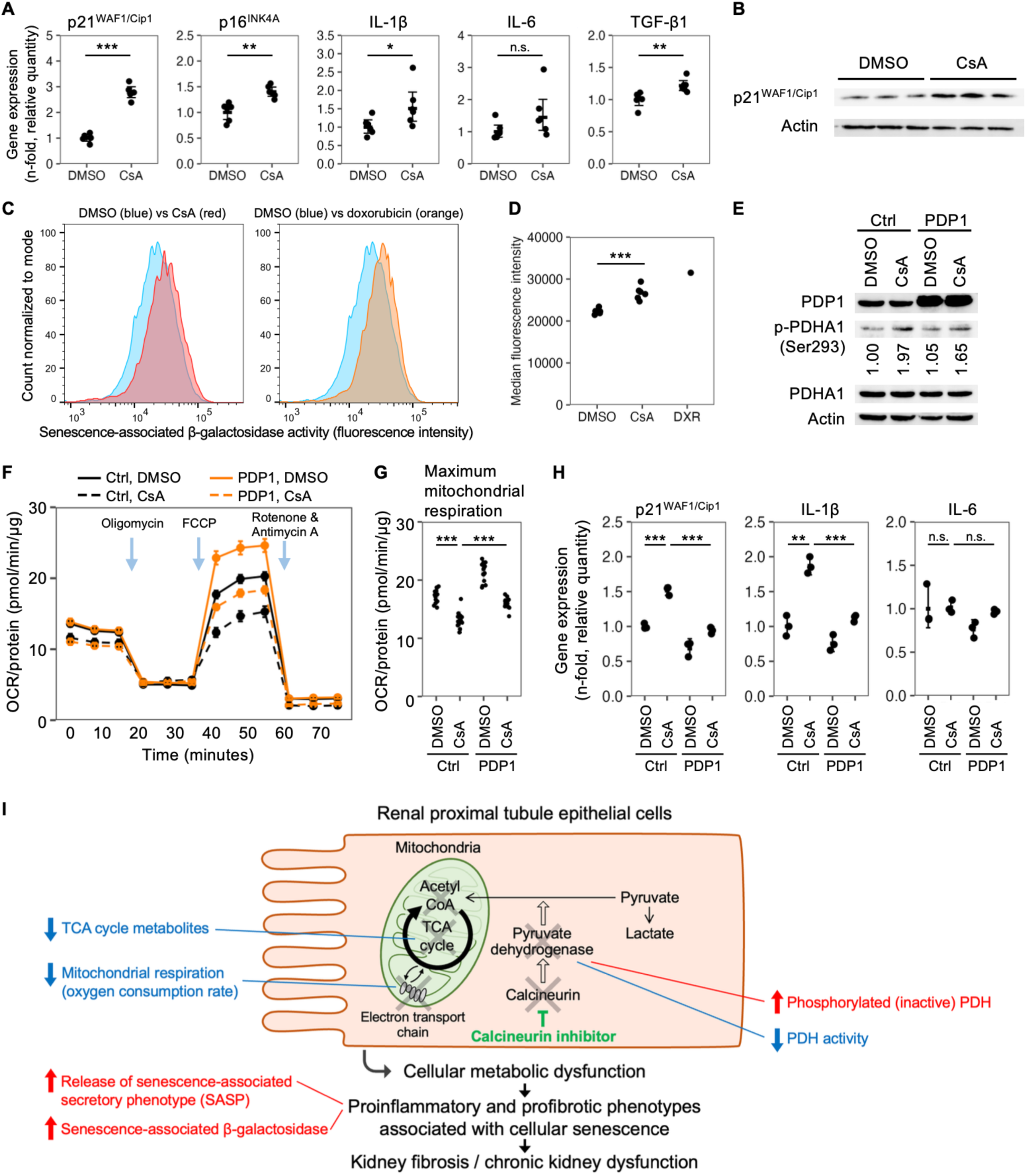
Activation of pyruvate dehydrogenase mitigates calcineurin inhibitor-induced metabolic dysfunction and cellular senescence–associated phenotypes. (A) mRNA expression of genes associated with cellular senescence measured by quantitative PCR in primary human renal proximal tubule epithelial cells (RPTECs) cultured with vehicle or cyclosporin A (CsA) for 24 h (n = 6 per group). (B) Western blot of p21^WAF1/Cip1^ and actin in primary human RPTECs cultured with DMSO or CsA for 5 days. (C and D) Flow cytometric assay for quantification of senescence-associated β-galactosidase activity in primary human RPTECs cultured with DMSO, CsA or doxorubicin for 24 h (C). Mean fluorescence intensity is plotted in (D). RPTECs cultured with doxorubicin were used as a positive control. (E) Western blot of PDP1, PDHA1 phosphorylation at Ser293, total PDHA1 and actin in primary human RPTECs transduced with lentiviral vectors encoding the gene of either a control protein (mCherry) or PDP1 and cultured with DMSO or CsA for 12 h. PDHA1, pyruvate dehydrogenase E1 subunit alpha 1; PDP1, pyruvate dehydrogenase phosphatase 1. (F and G) Oxygen consumption rate (OCR) normalized to total protein levels in primary human RPTECs transduced with lentiviral vectors encoding the gene of either a control protein (mCherry) or PDP1 and cultured with DMSO or CsA for 24 h (F). Maximum mitochondrial respiration is calculated and plotted in (G). n = 15 per group. (H) mRNA expression of genes associated with cellular senescence measured by quantitative PCR in primary human RPTECs transduced with lentiviral vectors encoding the gene of either a control protein (mCherry) or PDP1 and cultured with DMSO or CsA for 24 h (n = 3 per group). (I) Schematic illustration of proposed mechanism of chronic calcineurin inhibitor nephrotoxicity. Inhibition of calcineurin causes deactivation of pyruvate dehydrogenase. This results in a decreased flux of TCA cycle metabolites and decreased mitochondrial respiration. Cellular metabolic dysfunction causes proinflammatory, profibrotic phenotypes associated with cellular senescence and induce kidney fibrosis. Data are presented as mean ± 1.96 SE (A, F and H) or mean ± SD (D and G). Unpaired two-tailed Welch’s t test was used. Holm’s correction for multiple comparisons was used in (G and H). n represents the number of biological replicates. * *p* < 0.05, ** *p* < 0.01, *** *p* < 0.001. n.s., not significant. See also Figure S4.

### Activation of pyruvate dehydrogenase mitigates CNI-induced metabolic dysfunction and increase in SASP factor gene expression

To assess whether activation of pyruvate dehydrogenase mitigates metabolic dysfunction and senescence-associated phenotypes caused by CNI, primary human RPTECs were transduced with lentiviral vectors encoding either a control protein (mCherry) or pyruvate dehydrogenase phosphatase 1 (PDP1), which is an endogenous phosphatase and hence an activator of pyruvate dehydrogenase (Figure 5E). Extracellular flux analysis showed that overexpression of PDP1 ameliorated the CsA-induced decrease in maximal mitochondrial respiration in primary human RPTECs (Figures 5F and 5G). Moreover, PDP1 overexpression also reduced the CsA-induced increase in the gene expression of *CDKN1A* and *IL1B* (Figure 5H). These findings show that activation of pyruvate dehydrogenase mitigates metabolic dysfunction as well as the increase in SASP factor gene expression caused by CNI. Findings from these *in vitro* experiments are summarized in Figure 5I.

### DCA administration mitigates kidney fibrosis in chronic CNI nephrotoxicity mouse model

Finally, we examined whether activation of pyruvate dehydrogenase mitigates kidney fibrosis in the chronic CNI nephrotoxicity mouse model. DCA activates pyruvate dehydrogenase by inhibiting pyruvate dehydrogenase kinase, which phosphorylates and hence deactivates pyruvate dehydrogenase.^26^

Mice were offered *ad libitum* access to water or DCA-mixed water alongside daily subcutaneous injections of CsA or its vehicle for 4 weeks. The levels of creatinine and urea nitrogen in the plasma at the end of the experiment are shown in Figure 6B and Table S1. Average water consumption per day was 2.4–5.0 mL per mouse and tended to be lower in mice that received DCA-mixed water (Figure S5A); however, this did not result in an increase in plasma urea nitrogen levels (Figure 6B and Table S1), suggesting that the decrease in water consumption did not cause dehydration. Histological analysis of mouse kidneys demonstrated that DCA administration reduced the immunohistochemical staining for type I collagen (Figures 6C and 6D) as well as Sirius red staining (Figures 6E and 6F) in the chronic CNI nephrotoxicity mouse model. These changes were accompanied by a lower expression of the *Col1a1* gene (coding for the α1 chain of type I collagen) in the chronic CNI nephrotoxicity model mice that received DCA compared to the model mice that received water (Figure 6G). We also examined the expression of genes associated with proximal tubule injury, cellular senescence and fibrosis (Figure 6G and S5B). DCA administration also mitigated the expression of *Vcam1* (coding for VCAM-1), *Havcr1* (coding for KIM-1) and *Cdkn2a* (tested for mRNA variants coding for p16^INK4A^) in the chronic CNI nephrotoxicity mouse model (Figure 6G). These results demonstrate that DCA administration mitigates kidney fibrosis in the chronic CNI nephrotoxicity mouse model, possibly by counteracting CNI-induced proximal tubule injury and cellular senescence.

**Figure 6.**
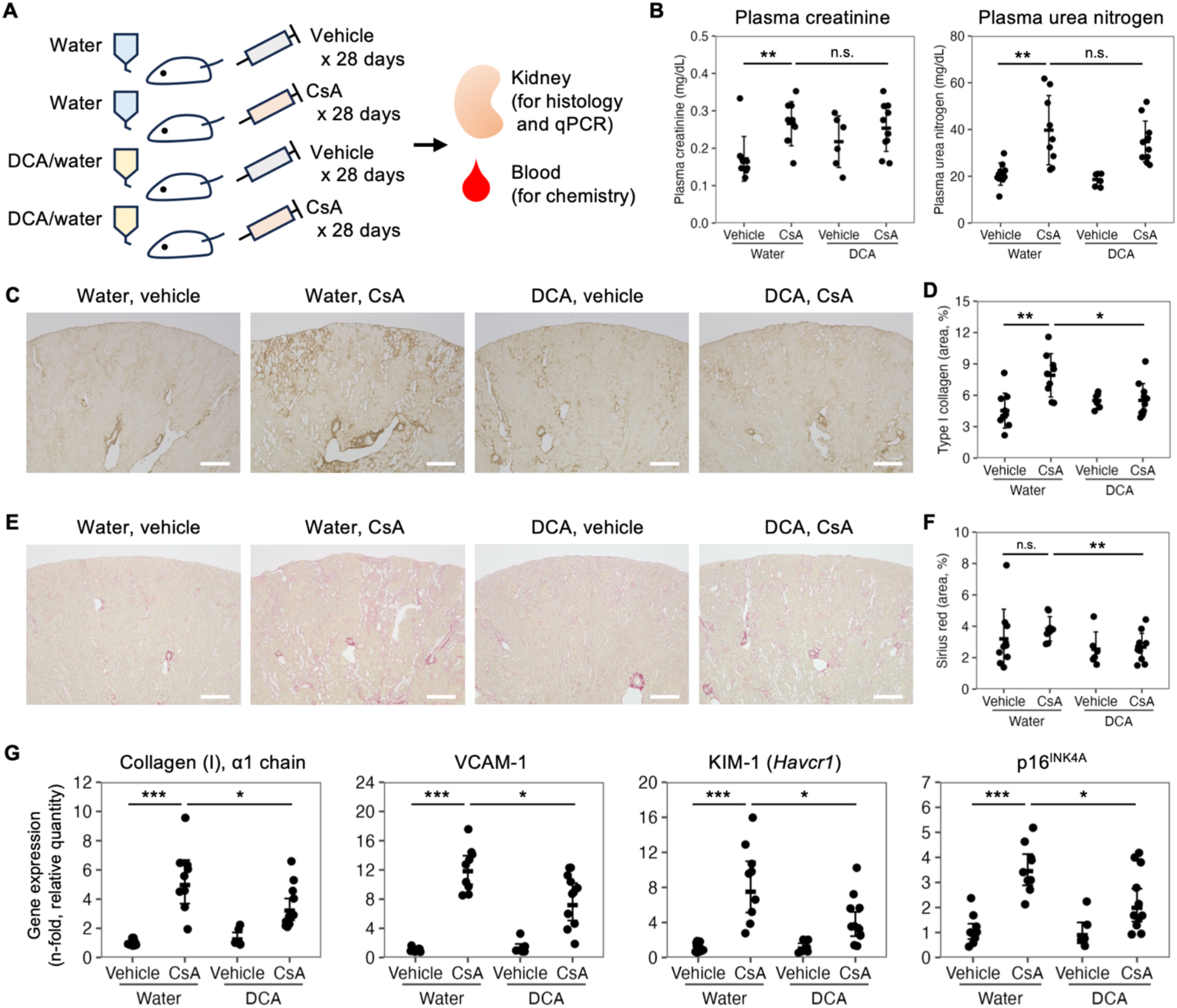
Dichloroacetic acid administration mitigates kidney fibrosis in chronic calcineurin inhibitor nephrotoxicity mouse model. (A) Experimental design of the experiment. CsA, cyclosporin A. (B) Plasma creatinine and urea nitrogen levels in ICR mice receiving vehicle or CsA as well as water or water mixed with dichloroacetic acid (DCA) for 4 weeks (n = 6–11 per group). (C and D) Immunostaining of type I collagen on kidney from ICR mice receiving vehicle or CsA as well as water or water mixed with dichloroacetic acid (DCA) for 4 weeks (D). Quantification of area percentage of immunostaining is plotted in (D). Scale bar, 500 μm. n = 6–11 per group. (E and F) Sirius red staining on kidney from ICR mice receiving vehicle or CsA as well as water or DCA-mixed water for 4 weeks (E). Quantification of area percentage of Sirius red staining is plotted in (F). Scale bar, 500 μm. n = 6–11 per group. (G) mRNA expression of genes associated with fibrosis, proximal tubule injury and cellular senescence measured by quantitative PCR in the kidneys of ICR mice receiving vehicle or CsA as well as water or DCA-mixed water (n = 6–11 per group). Data are presented as mean ± SD (B, D and F) or mean ± 1.96 SE (G). Unpaired two-tailed Welch’s t test with Holm’s correction for multiple comparisons was used. n represents the number of biological replicates. * *p* < 0.05, ** *p* < 0.01, *** *p* < 0.001. n.s., not significant. See also Table S1 and Figure S5.

## DISCUSSION

In this study, we elucidated the molecular mechanism of chronic CNI nephrotoxicity by focusing on the physiological role of calcineurin in regulating cellular metabolism in normal kidneys and how its inhibition can cause kidney fibrosis. Specifically, we found that calcineurin inhibition deactivates pyruvate dehydrogenase and induces proximal tubule cell metabolic dysfunction, causing profibrotic phenotype. These results provide a novel dimension to a fundamental understanding of the etiology of chronic CNI nephrotoxicity, where endothelial injury has long been the focus.^3,4^

Proximal tubule cells consume a large amount of energy for sodium reabsorption via the active transport of sodium through Na-K-adenosine triphosphatase in the basolateral membranes,^27^ making these cells vulnerable to metabolic dysfunction. Injury to proximal tubule cells alone can cause an inflammatory response and induce kidney fibrosis.^28^ Previous studies have shown that improving cellular metabolism by increasing mitochondrial biogenesis mitigates phenotypes associated with kidney injury or fibrosis in experimental murine models and in cultured kidney cells.^29–31^ Our results are consistent with the recent findings that dysregulated mitochondrial metabolism in proximal tubule cells can cause kidney fibrosis and is a potential target for interventions to ameliorate kidney fibrosis.

Although arteriolopathy is often found in chronic CNI nephrotoxicity and has been presumed to contribute to its etiology, a comprehensive analysis of the phenotypic changes in each cell type in chronic CNI nephrotoxicity had been lacking.^3,4^ We used snRNA-seq and found a significant increase in the proportion of KIM-1–positive injured proximal tubule cells with distinct transcriptional changes in the kidney of an early-stage chronic CNI nephrotoxicity mouse model. Together with our findings *in vitro* and *in vivo*, these results highlight how proximal tubule cells contribute to kidney fibrosis in chronic CNI nephrotoxicity.

The finding that CNIs impair mitochondrial energy metabolism is in line with a previous observation that human RPTECs cultured with CNIs show a decrease in mitochondrial respiration and in the expression of electron transport chain proteins.^25^ While the molecular mechanism of mitochondrial dysfunction has remained unknown, the present study shows that CNIs decrease the activity of pyruvate dehydrogenase and that activating pyruvate dehydrogenase mitigates the CNI-induced proinflammatory and profibrotic phenotypes. The idea that pyruvate dehydrogenase activity may be affected by calcineurin inhibition in proximal tubule cells stems from a recent finding that calcineurin directly dephosphorylates and thereby activates pyruvate dehydrogenase in an immortalized human cell line.^11^ We demonstrated that calcineurin inhibition results in increased phosphorylation of pyruvate dehydrogenase and hence decreased activity of pyruvate dehydrogenase in primary human RPTECs, whose cellular metabolism may differ from that of immortalized cells.

We also showed that mitochondrial dysfunction caused by calcineurin inhibition induces proinflammatory, profibrotic phenotypes associated with cellular senescence. This is consistent with a previous observation of the *CDKN1A* (coding for p21^WAF1/Cip1^) gene expression in cultured human proximal tubule cells.^32^ However, the expression of SASP factors had not been examined or discussed in previous studies, and whether the phenotypes associated with CNI-induced cellular senescence are associated with kidney fibrosis remained unknown. The present study revealed that CNIs induce proinflammatory, profibrotic phenotypes associated with cellular senescence not only *in vitro* but also *in vivo* and that these phenotypes are mitigated by pyruvate dehydrogenase activation. These findings are consistent with the fact that mitochondrial dysfunction is a major cause of cellular senescence.^19–21^

It is interesting that proinflammatory phenotypes are caused by calcineurin inhibition, which is generally associated with its immunosuppressive effects in T cells by down-regulating nuclear factor of activated T cells (NFAT) transcription factors. The upregulation of proinflammatory phenotypes by calcineurin inhibition *in vitro* and *in vivo* supports the idea that something other than NFAT downregulation takes place in the kidneys when CNIs are used.

The findings of this study suggest two theoretically possible approaches for chronic CNI nephrotoxicity. The first is to activate pyruvate dehydrogenase, preferably only in the kidneys. However, organ-specific activation of pyruvate dehydrogenase is not feasible with current technology. Second is to develop a novel immunosuppressant that suppresses NFAT but does not deactivate calcineurin.

In summary, the present study demonstrates that CNIs promote kidney fibrosis by deactivating pyruvate dehydrogenase and impairing proximal tubule cell energy metabolism. This study provides a novel perspective for understanding chronic CNI nephrotoxicity and paves the way for potential solutions.

### Limitations of the study

We were unable to use transgenic mice in this study because chronic CNI nephrotoxicity with kidney fibrosis develops in ICR mice^12^ but does so to a very limited extent in C57BL/6 mice and other strains. Therefore, proximal tubule–specific genetic interventions were not performed in this study. Instead, snRNA-seq and quantitative PCR of the gene coding for KIM-1 in the kidneys showed the involvement of injured proximal tubule cells in the disease etiology.

DCA administration was the only method used to activate pyruvate dehydrogenase *in vivo*. Although we sought to use multiple pyruvate dehydrogenase activators for *in vivo* experiments, we could not find any well-known established pyruvate dehydrogenase activators other than DCA. Developing and applying a novel DCA activator would extend these findings.

## METHODS

### Animal studies

All animal experiments were approved in advance by the Animal Care and Use Committee of the University of Tokyo Graduate School of Medicine (approval number A2023M092) and were performed in accordance with the Manual for Animal Experiment of the University of Tokyo. Jcl:ICR mice were purchased from CLEA Japan (Tokyo, Japan) and used for all animal experiments in this study.

The chronic calcineurin inhibitor mouse model was established as described in the literature^12^ with some modifications. Five-week-old male ICR mice weighing 28–33 g were housed in mouse cages in an air-, temperature-, and light-controlled environment. One week after the chow was switched to a low-sodium diet (0.01% sodium, Oriental Yeast, Tokyo, Japan), 30 mg/kg CsA (C2408, Tokyo Chemical Industry, Tokyo, Japan) solution or vehicle was subcutaneously administered to mice daily for 6 days, 2 weeks or 4 weeks, depending on the experiment. CsA was diluted in ethanol and then mixed with Kolliphor EL (#30554032, BASF, Ludwigshafen, Germany), which is a non-ionic solubilizer and emulsifier made by reacting castor oil with ethylene oxide. Kolliphor EL was previously called Cremophor EL and is a vehicle for commercial solution of CsA for human and animal use (Sandimmune Injection, Novartis, Basel, Switzerland). Food and drinking water were available *ad libitum*. Only male mice were used in this study because male rodents are more susceptible to chronic calcineurin inhibitor nephrotoxicity than female rodents.^38^

### Cell culture

Primary human RPTECs (CC-2553, Lonza, Walkersville, MD, USA) were used for in vitro studies instead of immortalized cell lines because immortalized cell lines could potentially have fundamental differences in cellular metabolism compared with primary cells. Primary human RPTECs were cultured in Epithelial Cell Medium (#4101, ScienCell Research Laboratories, Carlsbad, CA, USA) mixed with the following supplements provided by the manufacturer: 2% fetal bovine serum, 1% epithelial cell growth supplement, penicillin and streptomycin. Primary human RPTECs were cultured in a humidified incubator with 5% carbon dioxide at 37°C and were used for experiments within passages 2–5. Cell culture medium was renewed at least once every two days. Cells were washed with 4-(2-hydroxyethyl)-1-piperazineethanesulfonic acid (HEPES) buffered saline solution (CC-5024, Lonza) before renewing the medium or passaging the cells. Cell passaging was performed with 0.05% (w/v) trypsin solution (#204-16935, FUJIFILM Wako Pure Chemical Corporation, Osaka, Japan) and trypsin neutralizing solution (CC-5002, Lonza). To pellet the cells during the cell passaging, cell suspension was centrifuged at no more than 230 g for 3 minutes. CsA and tacrolimus (S5003, Selleck Chemicals, Houston, TX, USA) were the CNIs used in this study, and DMSO (#047-29353, FUJIFILM Wako Pure Chemical Corporation) was used as vehicle. The final concentration of DMSO in the culture medium was 0.1% (v/v) or below.

### Human data

Publicly available microarray datasets of human kidney allografts (GSE53605 and GSE178689) were analyzed with GEO2R, an analysis tool available on the Gene Expression Omnibus website (https://www.ncbi.nlm.nih.gov/geo/), to find differentially expressed genes (DEG) between sample groups within each dataset.

### Single-nucleus RNA sequencing (snRNA-seq)

Nuclear dissociation was performed using the Nuclei EZ Prep Kit (NUC101, Merck KGaA, Darmstadt, Germany) as described previously.^39^ Kidneys from one early-stage chronic calcineurin inhibitor nephrotoxicity model mouse after 6 days of cyclosporin A administration and from one control mouse after 6 days of vehicle administration were subjected to analysis. A thin, horizontally sliced, snap-frozen kidney was minced with a razor blade and homogenized in ice-cold EZ Lysis Buffer. After 5 min of incubation on ice, the homogenate was filtered through a 40-μm strainer (#43-50040, PluriSelect, El Cajon, CA, USA) and centrifuged at 500 g for 5 min. The pellet was resuspended with EZ Lysis Buffer, centrifuged at 500 g for 5 min, resuspended in Dulbecco’s phosphate buffered saline (PBS) with 1% fetal bovine serum and filtered through a 5-μm strainer (#43-50005, PluriSelect).

Formation of gel beads in emulsion, barcoding, cDNA amplification and library construction were performed according to the manufacturer’s instructions (PN-1000269, Chromium Next GEM Single Cell 3ʹ Kit v3.1, 10x Genomics, Pleasanton, CA, USA). cDNA libraries were sequenced on an Illumina NovaSeq instrument, generating 2 × 150 bp paired-end reads. Sequencing data were analyzed using Cell Ranger (10x Genomics) with reference to refdata-gex-mm10-2020-A dataset provided by 10x Genomics. Major quality control metrics are summarized in Table S2.

Downstream analysis was performed with Seurat package (version 5.0.3)^33^ in R system for statistical computing (version 4.4.0, R Foundation for Statistical Computing, Vienna, Austria). After removal of ambient RNA with SoupX (version 1.6.2),^34^ low-quality nuclei, defined by either mitochondrial transcript percentage of ≥ 1% or detected gene numbers of ≤ 200 or ≥ 7000, were filtered out. DoubletFinder (version 2.0.4)^35^ was used for doublet detection with an assumption that 5.6% of the nuclei were doublets. Following the removal of doublets detected at this point, data from early-stage CNI nephrotoxicity mouse model and its control were merged and integrated with IntegrateLayers function to perform anchor-based canonical correlation analysis integration.

Unsupervised clustering of 12,743 cells identified 18 cell clusters, which included a small cluster of 48 cells (0.38% of all cells) annotated as doublets with linage markers (Figure S1) that were not detected by DoubletFinder. These doublets were removed, and the remaining 12,695 cells in 17 clusters encompassing all major cell types in the kidneys were subjected to analysis.

### Gene Set Enrichment Analysis (GSEA)

GSEA of snRNA-seq dataset at single-cell resolution was performed with the VISION package (version 3.0.1).^36^ Genes associated with oxidative phosphorylation, glycolysis and TGF-β signaling were analyzed with hallmark gene sets with reference to mh.all.v2023.2.Mm.symbols.gmt file available on the GSEA website. Genes related with cellular senescence were analyzed with Gene Ontology gene sets with reference to m5.all.v2023.2.Mm.symbols.gmt file available on the GSEA website, because hallmark gene sets do not contain a list of genes associated with cellular senescence.

Kyoto Encyclopedia of Genes and Genomes (KEGG) pathway enrichment analysis was performed for DEG between two cell clusters or two groups of human kidney allograft with WEB-based GEne SeT AnaLysis Toolkit (WebGestalt).^37^ Genes were ranked based on log FC value. Figures 2A, 2B, 3A and 3B were generated by WebGestalt.

### Histological analysis

Formalin-fixed, paraffin-embedded kidney sections were cut at 3 μm thickness and subjected to Masson’s trichrome staining, Sirius red staining and immunostaining for assessment of kidney fibrosis.

Immunohistochemical staining of type I collagen on mouse kidney sections was performed as follows. Sections were deparaffinized in Histo-Clear (HS-200, National Diagnostics, Atlanta, GA, USA), rehydrated in ethanol and washed with PBS. Heat-induced epitope retrieval was performed by soaking the sections in boiling citrate buffer for 10 minutes. Sections were allowed to cool at room temperature for at least 30 minutes, washed with PBS, blocked with Protein Block (X090930, Agilent Technologies, Santa Clara, CA, USA) and incubated with goat anti-type I collagen polyclonal antibody (1:100, #1310-01, SouthernBiotech, Birmingham, AL, USA) overnight at 4℃. Sections were then washed with PBS, incubated with Histofine Simple Stain Mouse MAX-PO G (#414351, Nichirei Biosciences, Tokyo, Japan), washed with PBS and visualized with ImmPACT DAB (SK-4105, Vector Laboratories, Newark, CA, USA), washed with water, dehydrated with ethanol, cleared with Histo-Clear and mounted with Mount-Quick (DM-01, Daido Sangyo, Tokyo, Japan).

Whole slide images were obtained with a BZ-X710 microscope (Keyence, Osaka, Japan). After trimming off the renal papilla and renal pelvis wall from the images, area of positive staining and the total area of kidney tissue were detected with ImageJ software (National Institutes of Health, Bethesda, MD, USA) by applying the same threshold value to all images.

### Quantitative PCR

RNA extraction from kidney tissue was performed by soaking a snap-frozen kidney tissue in RNAiso Plus (#9109, Takara Bio, Shiga, Japan) with 1.4-mm zirconium oxide beads (P000927-LYSK0-A.0, Bertin Technologies, Montigny-le-Bretonneux, France) and mechanically homogenizing the tissue with a Minilys homogenizer (Bertin Technologies). RNA extraction from culture cells was conducted by washing the cells with HEPES buffered saline solution and soaking the cells in RNAiso Plus (Takara Bio). Total RNA was isolated according to the manufacturer’s instructions. The concentration and quality of RNA were quantified with a BioSpec-nano spectrophotometer (Shimadzu Corporation, Kyoto, Japan). cDNA was created with PrimeScript RT Master Mix (RR036B, Takara Bio) and was subjected to real-time quantitative PCR with THUNDERBIRD SYBR qPCR Mix (QPS-201, Toyobo, Tokyo, Japan) and CFX Connect Real-Time PCR Detection System (Bio-Rad, Hercules, CA, USA). Primer sequences are described in Table S3. Gene expression was normalized to β-actin mRNA levels. Relative gene expression levels of two groups were compared by testing the ΔΔ quantification cycle (ΔΔCq) values.

### Western blot

Cells were suspended in radioimmunoprecipitation assay (RIPA) buffer (#188-02453, FUJIFILM Wako Pure Chemical Corporation) with Halt Protease and Phosphatase Inhibitor Single-Use Cocktail (#78442, Thermo Fisher Scientific, Waltham, MA, USA). After 15 minutes of incubation on ice and 10 minutes of centrifugation, supernatant was collected and stored at −80°C. Protein levels were quantified by Pierce BCA Protein Assay Kit (#23225, Thermo Fisher Scientific). Protein solution was mixed with sodium dodecyl sulfate (SDS) sample buffer, which consisted of 60 mM Tris-HCl (pH 6.8), 2% sodium dodecyl sulfate, 10% glycerol, 0.012% bromophenol blue and 0.01 M dithiothreitol, and were boiled at 95°C for 5 minutes. Proteins were separated with 8–12% SDS-polyacrylamide gel electrophoresis and transferred to polyvinylidene difluoride membranes (Amersham Hybond P 0.45 PVDF blotting membrane, #10600023, Cytiva, Marlborough, MA, USA) with transfer buffer by Trans-Blot Turbo Transfer System (Bio-Rad). Transfer buffer consisted of 48 mM Tris-base buffer, 39 mM glycine, 0.04% SDS and 20% methanol. To prevent nonspecific protein binding, membranes were shaken in Tris-buffered saline (pH 7.4) with 5% bovine serum albumin and 0.05% Tween 20 at room temperature for 30 minutes. Membranes were incubated with primary antibodies in Tris-buffered saline with 0.05% Tween 20 (TBS-T) for 1 h at room temperature or overnight at 4°C, washed with TBS-T three times for 5 minutes each, incubated with secondary antibodies in TBS-T for 45–60 minutes at room temperature, and washed with TBS-T four times for 5 minutes each. Blots were visualized by Pierce ECL Plus Western Blotting Substrate (#32132, Thermo Fisher Scientific) and ImageQuant LAS 4000 (FUJIFILM, Tokyo, Japan). Blots were stripped with Western BLoT Stripping Buffer (T7135A, Takara Bio) and reprobed when necessary. Densitometry analysis was performed with ImageJ software (National Institutes of Health). Antibodies used in this study are as follows: rabbit anti-phospho-PDH α1 (Ser293) polyclonal antibody (1:1,000; #31866, Cell Signaling, Danvers, MA, USA), mouse anti-PDHA1 monoclonal antibody (1:10,000; #66119-1-Ig, Proteintech Group, Rosemont, IL, USA), rabbit anti-p21 monoclonal antibody (1:1,000; ab109199, Abcam, Cambridge, UK), rabbit anti-PDP1 monoclonal antibody (1:1,000, #65575, Cell Signaling), rabbit anti-actin polyclonal antibody (1:2,000; A2066, Merck KGaA), goat anti-rabbit IgG (H + L)-HRP conjugate (1:10,000; #170-6515, Bio-Rad), goat anti-mouse IgG (H + L)-HRP conjugate (1:10,000; #170-6516, Bio-Rad).

### Extracellular flux analysis

Seahorse XFe96 Analyzer (Agilent Technologies, Santa Clara, CA, USA) was used for the measurement of OCR of primary human RPTECs seeded on 96-well XF96 Cell Culture Microplates (#101085-004, Agilent Technologies) at 5,000–7,000 cells per well depending on the experiment. After confirming that the cells were attached to the plates, culture medium was changed to a fresh medium with either 10 μM cyclosporin A, 10 μM tacrolimus or their vehicle. Twenty-four hours later, medium was changed to assay medium. Assay medium consisted of XF Base Medium Minimal DMEM with phenol red (#103334-100, Agilent Technologies), 1 mM pyruvate, 2 mM glutamine, 10 mM glucose and other chemicals depending on the experiment. Following a 45-minute incubation in a 37°C non-CO2 incubator, microplates were subjected for OCR measurement. Key parameters of mitochondrial respiration were measured by applying Seahorse XF Cell Mito Stress Test Kit (#103015-100, Agilent Technologies), which is a set of oligomycin, carbonyl cyanide 4-(trifluoromethoxy)phenylhydrazone (FCCP) and a mixture of rotenone and antimycin A. Following the measurement of basal respiration, oligomycin was added to the assay medium in the microplate to inhibit ATP synthase, also known as complex V. After three measurements of OCR, FCCP was added to the assay medium to collapse the mitochondrial proton gradient and induce maximal respiration. Finally, rotenone and antimycin A were added to the assay medium to measure OCR when mitochondrial respiration is halted by a complete block of electron transportation chain. OCR associated with maximal respiration was calculated by subtracting non-mitochondrial OCR (measured after adding rotenone and antimycin A) from maximal OCR (measured after adding FCCP).

After the OCR measurement was completed, assay medium was removed from the microplates, and radioimmunoprecipitation assay (RIPA) buffer was applied to each well to extract proteins from the cells. The amount of protein in each well was determined by BCA assay. The OCR was normalized to the amount of protein, which reflects the number of cells, in order to remove the effect of potential changes in OCR caused by possible differences in cell numbers.

### Biochemical analysis

Plasma creatinine and urea nitrogen levels were evaluated with commercial kits (#469-07594 and #465-07694 for creatinine assay; #466-54891 and #462-54991 for urea nitrogen assay; all products from FUJIFILM Wako Pure Chemical Corporation). Pyruvate dehydrogenase activity was measured with Pyruvate Dehydrogenase Activity Assay Kit (MAK183, Merck KGaA). Protein concentration was determined by BCA assay. Absorbance was measured with EnSpire Multimode Plate Reader (PerkinElmer, Waltham, MA, USA)

### Gas chromatography–mass spectrometry (GC–MS)

RPTECs were subjected to GC–MS to analyze water-soluble metabolites related to TCA cycle. Water-soluble metabolites were extracted from primary human RPTECs as described previously^40^ with some modifications. RPTECs on 10-cm dishes were washed twice with ice-cold HEPES buffered saline solution and were covered with a mixed solution of 800 μL methanol of liquid chromatography–mass spectrometry grade (#134-14523, FUJIFILM Wako Pure Chemical Corporation) and 5 μL of 1 mg/mL 2-isopropylmalic acid, which served as an internal standard. Cell debris and the solution were collected to 15-mL conical centrifuge tubes with scrapers and pipette tips, snap frozen in liquid nitrogen, and stored at −80°C. On a subsequent day, samples were taken out from the freezer and added with 800 μL chloroform cooled in a −30°C freezer followed by 320 μL double distilled water (DDW) cooled to 4°C. The solutions were vortexed, sonicated with an ultrasonic bath sonicator for 60 seconds, vortexed again for 60 seconds, and centrifuged at 2,900 g for 20 minutes. The supernatant was transferred to new 15-mL centrifuge tubes, vortexed, and evaporated to dryness at 40°C in a vacuum evaporator. Standard mixtures of metabolites for GC–MS (#1021-58400, GL Sciences, Tokyo, Japan) were also evaporated and served as a reference for metabolite identification.

Dried samples were subjected to derivatization for GC–MS analysis. Samples were rigorously mixed with 80 μL for samples or 40 μL for standards of 40 mg/mL methoxyamine hydrochloride (#226904, Merck KGaA) in pyridine (#270970, Merck KGaA) in a fume hood, centrifuged at 2,900 g for 2 minutes, mixed by pipetting, and incubated in a 37°C water bath for 90 minutes. Samples were added with 160 μL of N-methyl-N-trimethylsilyl-trifluoroacetamide (#1022-11060, GL Sciences) in a fume hood, centrifuged at 2,900 g for 2 minutes, mixed by pipetting, and incubated in a 37°C water bath for 30 minutes. Samples were stored at −80°C for more than 72 hours before subjected to GC–MS analysis to stabilize the derivatization status.

Metabolite analysis was performed with a GCMS-QP2010 Plus (Shimadzu Corporation) gas chromatograph mass spectrometer, operating in full scan mode, equipped with a DB-5 column (inner diameter, 0.25 mm; length, 30 m; film thickness, 1 μm; #122-5033, Agilent Technologies). Helium was used as carrier gas. An aliquot of derivatized sample was injected to a GC-2010 gas chromatograph at a split ratio of 1:10. Vaporization chamber temperature was set at 280°C. The column oven temperature was initially held at 100°C for four minutes, and then linearly increased at the rate of 10°C per minute until it reached 320°C, where it was kept for 11 minutes. Ion source temperature was 200°C, and the GC–MS interface temperature was 280°C. The data were analyzed with GCMSsolution software version 4.52 (Shimadzu Corporation). In addition to mass spectra derived from the standard mixtures, Smart Metabolites Database version 2 (Shimadzu Corporation) served as the reference for spectrum analysis. All mass spectra were subject to automatic detection as well as manual inspection followed by manual correction if necessary. The area under the curve (AUC) was measured for one representative derivative of each metabolite. In order to remove the effect of inter-sample difference in the efficiency of metabolite extraction and derivatization, the AUCs of all representative derivatives were divided by the AUC of a representative derivative of 2-isopropylmalic acid in each sample.

### Senescence-associated β-galactosidase activity assay with flow cytometry

Senescence-associated β-galactosidase activity was detected using the Cellular Senescence Detection Kit - SPiDER-βGal (SG03, Dojindo Laboratories, Kumamoto, Japan). Cells were stained with the kit according to the manufacturer’s instructions, washed with Hank’s balanced salt solution (HBSS) twice, and collected with a scraper. After cells were pelleted with 3 minutes of centrifugation at 500 g, cells were resuspended in HBSS and subjected to flow cytometry with CytoFLEX (Beckman Coulter, Brea, CA, USA). FlowJo software (BD, Franklin Lakes, NJ, USA) was used for data analysis.

### Lentiviral vector packaging and transduction

Lentiviral vectors that encoded PDP1 or a control protein (mCherry) were purchased from VectorBuilder Japan (Kanagawa, Japan). Both vectors encoded enhanced green fluorescent protein (EGFP) and puromycin resistance gene. Primary human RPTECs were incubated with culture medium added with lentiviral vectors and 8 µg/mL polybrene for 8 hours, after which the medium was renewed. Two days later, culture medium was changed to fresh medium with 3.5 µM puromycin, which was lethal to most normal primary human RPTECs after 2 days of incubation. Two days later, culture medium was renewed, and the fluorescence of EGFP was confirmed with BZ-X710 (Keyence) fluorescence microscope.

All experiments were approved in advance by The University of Tokyo Graduate School of Medicine Committee for Living Modified Organisms (approval number 25-8) and were conducted in accordance with The University of Tokyo Rules for the Use of Living Modified Organisms.

### Statistical analysis

Data were analyzed using R system for statistical computing (version 4.4.0, R Foundation for Statistical Computing) unless otherwise noted. Unpaired two-tailed Welch’s t test was performed to compare the mean of two groups. The level of statistical significance was set at *p* < 0.05. When multiple comparisons were made, significance level was adjusted following the Holm’s method.

## Acknowledgements

This work was supported by JSPS KAKENHI (Grant Number 23KJ0437 to Y.O., 24K11425 to H.N., and 23H02924 to M.N.) and the NIDDK Intramural Research Program (NIDDK, NIH, Bethesda, MD, USA). Y.O. acknowledges support from JST SPRING Grant Number JPMJSP2108. Y.O. is a Research Fellow of Japan Society for the Promotion of Science. The authors thank Ms. Harumi Yamamura at The University of Tokyo Graduate School of Medicine for her technical support in histological staining.

## Author contributions

Y.O. and H.N. conceptualized the study. Y.O. performed all the animal studies, cell culture experiments, data curation, and formal analysis. F.H. and Y.O. performed gas chromatography– mass spectrometry. T.Y. and Y.O. performed single-nucleus RNA sequencing. Y.O. wrote the original manuscript. H.N. and J.B.K. reviewed and edited the manuscript. Y.K, J.B.K and M.N. provided supervision. All authors read and approved the final manuscript.

## Supplemental information

Tables S1–S3 and Figures S1–S5

