## Supplementary material for "Calcineurin inhibition deactivates pyruvate dehydrogenase and induces proximal tubule cell metabolic dysfunction, causing profibrotic phenotype": Tables S1-S3 and Figures S1-S5

**Table S1. Water consumption of mice that received water or dichloroacetic acid (DCA) alongside vehicle or cyclosporin A, related to Figure 6B**

| Drink | Water | Water | DCA | DCA |
|---|---|---|---|---|
| Subcutaneous injection | Vehicle | CsA | Vehicle | CsA |
| Plasma creatinine level (mg/dL) | 0.17 ± 0.06 | 0.27 ± 0.06 | 0.22 ± 0.07 | 0.25 ± 0.06 |
| Plasma urea nitrogen level (mg/dL) | 20.9 ± 4.8 | 39.7 ± 14.8 | 18.6 ± 2.8 | 34.9 ± 8.7 |

* Data are presented as mean ± SD.

**Table S2. Quality control metrics for single-nucleus RNA sequencing**

| Item | | Vehicle | Cyclosporin A |
|---|---|---|---|
| Sequencing | Total RNA reads | 503,783,200 | 506,058,208 |
| | Valid barcodes | 95.6% | 96.4% |
| | Valid UMIs | 99.9% | 99.9% |
| | Sequencing saturation | 75.3% | 67.9% |
| Mapping | Reads mapped to genome | 90.1% | 92.7% |
| | Reads mapped confidently to genome | 88.2% | 91.0% |
| Cells | Estimated number of cells | 5,894 | 7,612 |
| | Mean reads per cell | 85,474 | 66,482 |
| | Median genes per cell | 2,336 | 2,444 |
| | Total genes detected | 24,545 | 24,443 |

**Table S3. Primer sequence for quantitative PCR**

| Species | Gene | Forward | Reverse |
|---|---|---|---|
| Mouse | *Actb* | AAGATCAAGATCATTGCTCCTCCTG | AAACGCAGCTCAGTAACAGTCC |
| | *Cdkn1a* | CGGTGTCAGAGTCTAGGGGA | AGGATTGGACATGGTGCCTG |
| | *Cdkn2a* | CTTTGTGTACCGCTGGGAAC | GCCGGATTTAGCTCTGCTCT |
| | *Col1a1* | GACGCATGGCCAAGAAGACA | CCTTGGGTCCCTCGACTCC |
| | *Col3a1* | AGCCTTCTACACCTGCTCCT | CGGATAGCCACCCATTCCTC |
| | *Havcr1* | AAACCAGAGATTCCCACACG | GTCGTGGGTCTTCCTGTAGC |
| | *Il6* | TCCAGTTGCCTTCTTGGGAC | GTGTAATTAAGCCTCCGACTTG |
| | *Tgfb1* | GCAACAATTCCTGGCGTTACC | CGAAAGCCCTGTATTCCGTCT |
| | *Vcam1* | ATGTCAACGTTGCCCCCAAG | AATGCCGGAATCGTCCCTTT |
| Human | *ACTB* | TCCCCCAACTTGAGATGTATGAAG | AACTGGTCTCAAGTCAGTGTACAGG |
| | *CDKN1A* | CTGCCTTAGTCTCAGTTTGTGT | AACCTCTCATTCAACCGCCTA |
| | *CDKN2A* | GGGTCGGGTAGAGGAGGTG | GCTGCCCATCATCATGACCT |
| | *IL1B* | CCACGGCCACATTTGGTT | AGGGAAGCGGTTGCTCATC |
| | *IL6* | ACTCACCTCTTCAGAACGAATTG | CCATCTTTGGAAGGTTCAGGTTG |
| | *TGFB1* | TACCTGAACCCGTGTTGCTCTC | GTTGCTGAGGTATCGCCAGGAA |

* The primers for *Cdkn2a* detect a messenger RNA that encodes $p16^{INK4A}$ but not $p19^{ARF}$.
The primers for *CDKN2A* detect a messenger RNA that encodes $p16^{INK4A}$ but not $p14^{ARF}$.

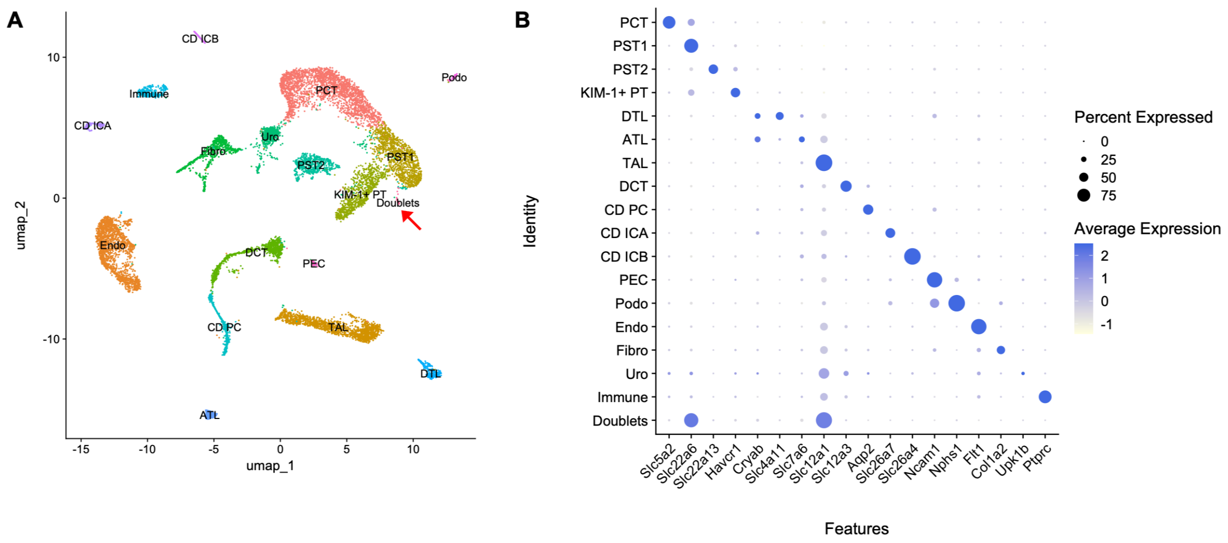


**Figure S1. Single-nucleus transcriptional profiling**

(A) UMAP presentation of 12,743 cells profiled from the kidney of early-stage chronic calcineurin inhibitor nephrotoxicity mouse model and its control. Red arrow designates a small cluster of 48 cells (0.38% of all cells) annotated as Doublets.

(B) Dot plot showing the expression pattern of cluster-specific marker genes.

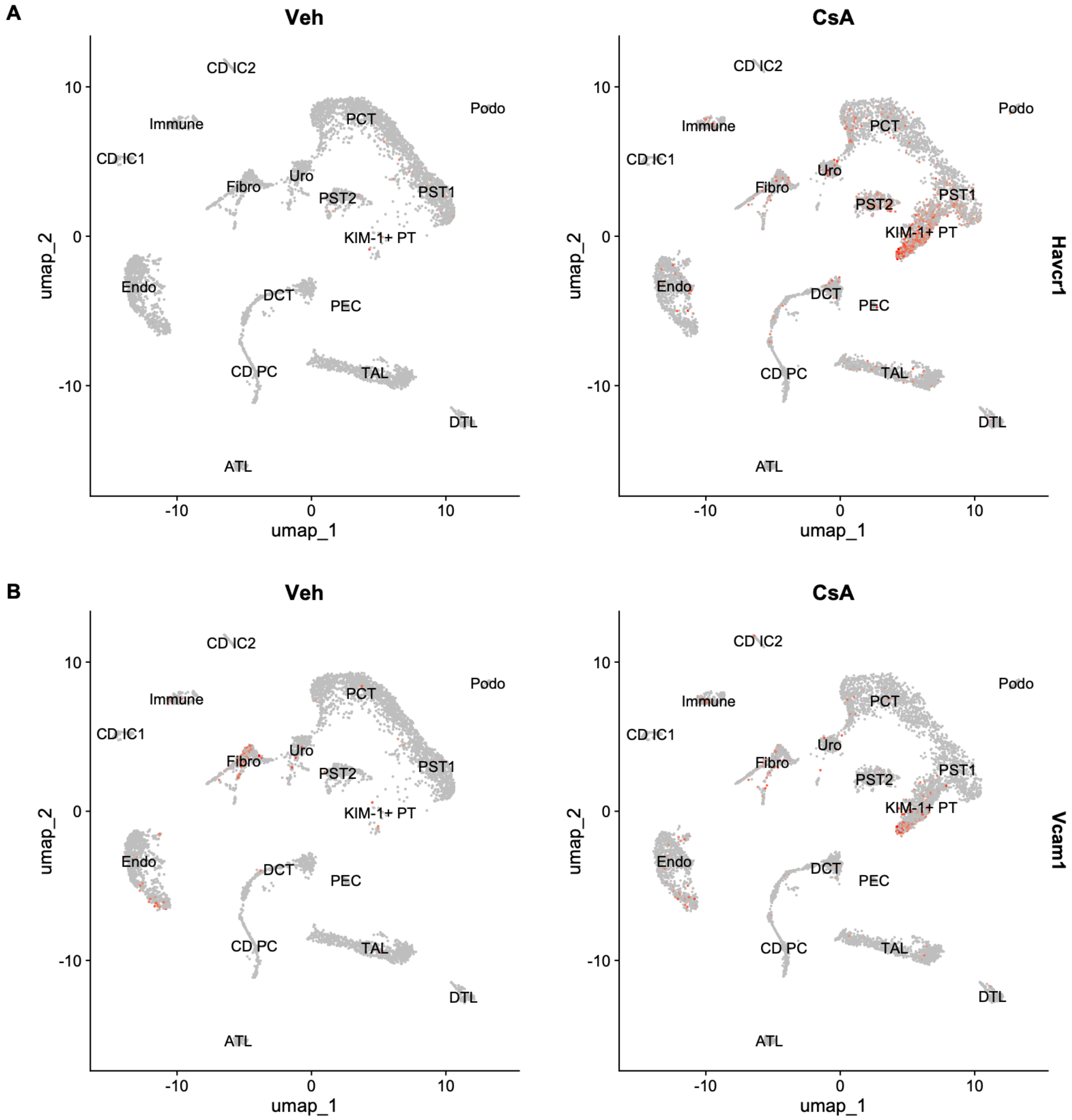


**Figure S2. UMAP presentation of the expression of genes *Havcr1* and *Vcam1* in the kidney of early-stage chronic calcineurin inhibitor nephrotoxicity mouse model and its control**
(A) Feature plot showing the expression of gene *Havcr1* (coding for KIM-1).
(B) Feature plot showing the expression of gene *Vcam1*.

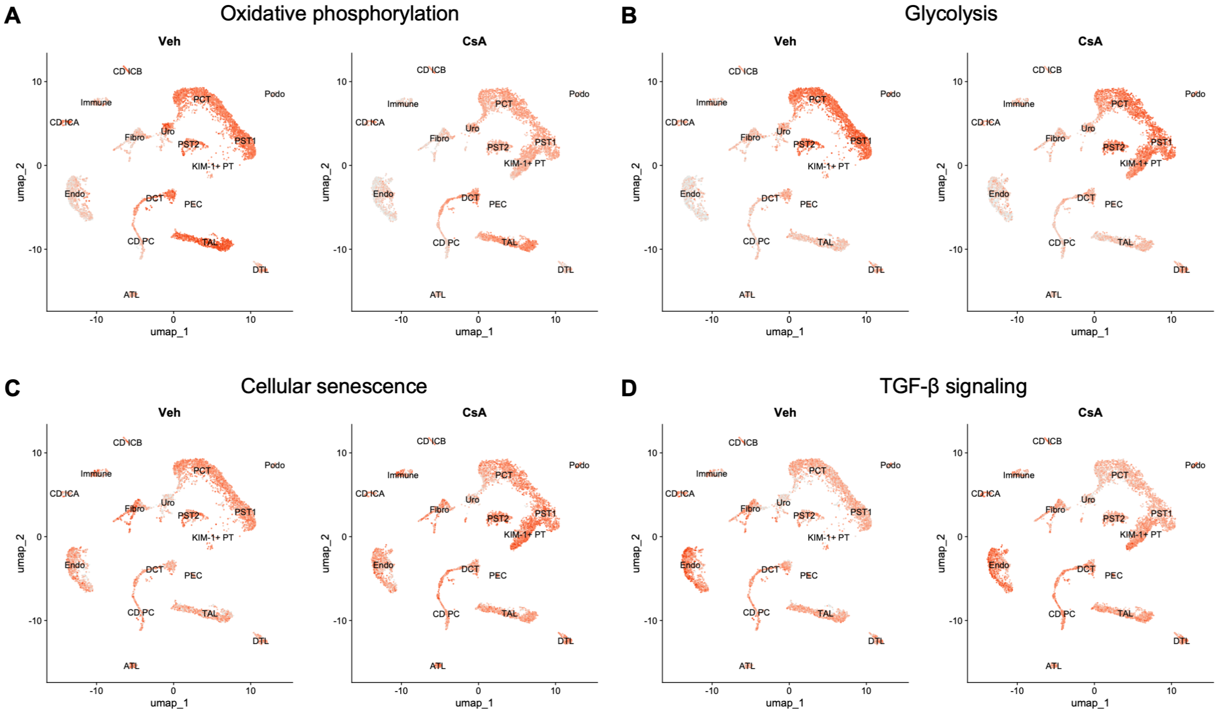


**Figure S3. UMAP displaying enrichment of genes associated with metabolic pathways, cellular senescence and fibrosis**

(A–D) Feature plot showing the enrichment of genes associated with oxidative phosphorylation (A), glycolysis (B), cellular senescence (C) and TGF-β signaling (D).

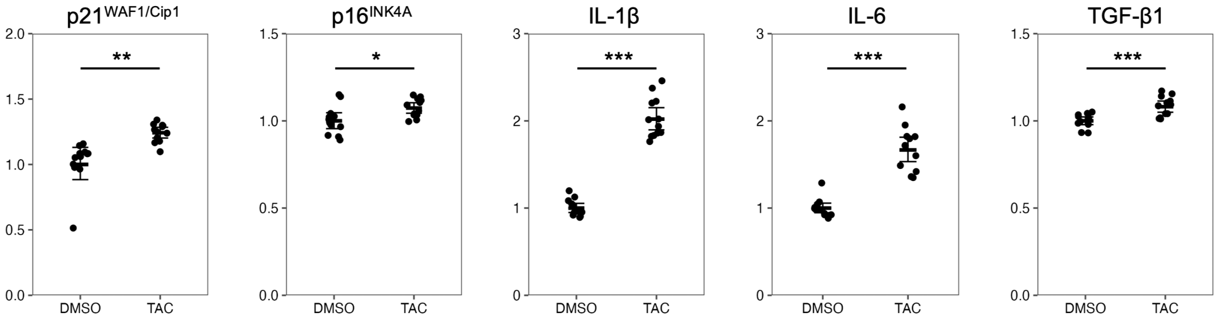


**Figure S4. Increased expression of genes associated with cellular senescence caused by tacrolimus**

mRNA expression of genes associated with cellular senescence measured by quantitative PCR in primary human renal proximal tubule epithelial cells cultured with vehicle or tacrolimus (TAC) for 24 h (n = 12 per group). Data are presented as mean ±1.96 SE. Unpaired two-tailed Welch's t test was used. n represents the number of biological replicates. * $p < 0.05$, ** $p < 0.01$, *** $p < 0.001$.

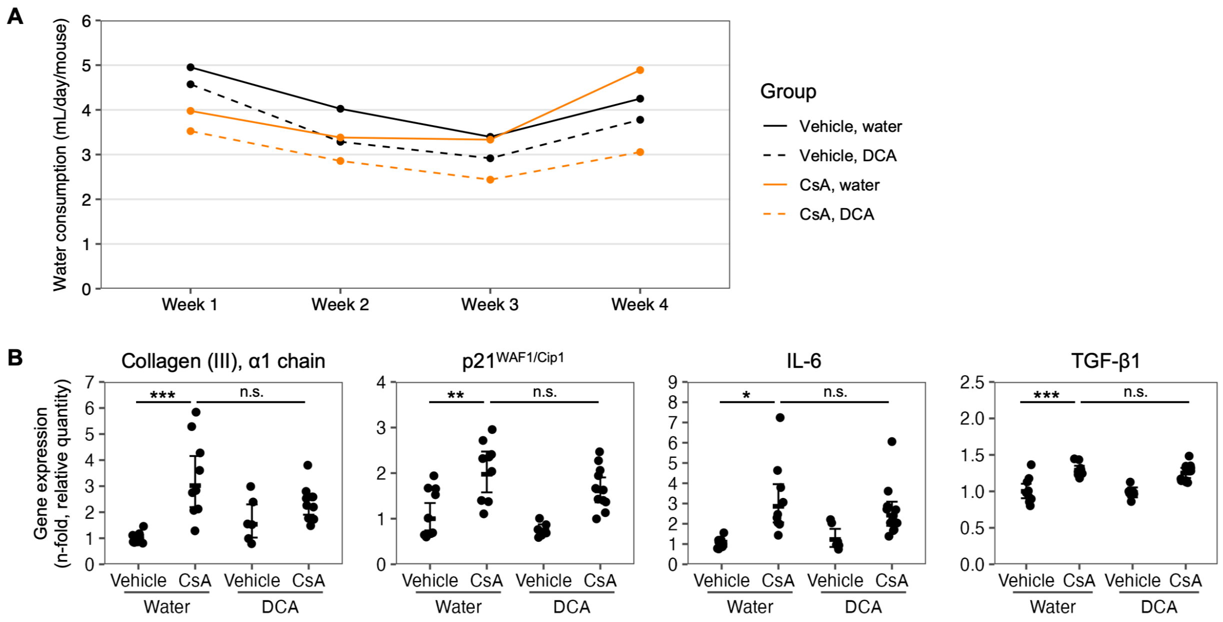


**Figure S5. Trajectory of water consumption and additional results on gene expression in the kidney**

(A) Trajectory of water consumption of mice. Values were calculated by dividing the total water consumption per cage per week by the number of mice in the cage and by the number of days per week.

(B) mRNA expression of genes associated with fibrosis and cellular senescence measured by quantitative PCR in the kidneys of ICR mice receiving vehicle or CsA as well as water or DCA-mixed water (n = 6–11 per group) in addition to Figure 6G. Data are presented as mean ± 1.96 SE. Unpaired two-tailed Welch's t test with Holm's correction for multiple comparisons was used. n represents the number of biological replicates. * $p < 0.05$, ** $p < 0.01$, *** $p < 0.001$. n.s., not significant.
